# Extracellular matrix micropatterning technology for whole cell cryogenic electron microscopy studies

**DOI:** 10.1101/657072

**Authors:** Leeya Engel, Guido Gaietta, Liam P. Dow, Mark F. Swift, Gaspard Pardon, Niels Volkmann, William I. Weis, Dorit Hanein, Beth L. Pruitt

**Affiliations:** Department of Bioengineering, Stanford University, Stanford, California; Immunity and Pathogenesis Program, Sanford Burnham Prebys Medical Discovery Institute, La Jolla, California; Biomolecular Science and Engineering Program, University of California, Santa Barbara; Departments of Mechanical Engineering and Molecular, Cellular, and Developmental Biology, University of California, Santa Barbara; Departments of Structural Biology and Molecular Cellular Physiology, Stanford University School of Medicine

## Abstract

Cryogenic electron tomography is the highest resolution tool available for structural analysis of macromolecular organization inside cells. Micropatterning of extracellular matrix (ECM) proteins is an established *in vitro* cell culture technique used to control cell shape. Recent traction force microscopy studies have shown correlation between cell morphology and the regulation of force transmission. However, it remains unknown how cells sustain increased strain energy states and localized stresses at the supramolecular level. Here, we report a technology to enable direct observation of mesoscale organization in epithelial cells under morphological modulation, using a maskless protein photopatterning method to confine cells to ECM micropatterns on electron microscopy substrates. These micropatterned cell culture substrates can be used in mechanobiology research to correlate changes in nanometer-scale organization at cell-cell and cell-ECM contacts to strain energy states and traction stress distribution in the cell.

## INTRODUCTION

To maintain epithelial homeostasis, cells must actively respond to static and dynamic mechanical cues from the ECM and neighboring cells. The size and structure of the cell microenvironment are crucial regulators of cell architecture, mechanics, polarity and function (1, 2). Micropatterning of cell culture substrates is increasingly being used as a tool to modify the structure of the cell microenvironment for *in vitro* studies. The geometry of micropatterned substrates has been used in conjunction with light microscopy to modulate the establishment of cell polarity and cell growth (3), cell viability (4), terminal differentiation (5), stem cell differentiation (6, 7), and cell division axis orientation (8). Tee et al. used correlated light microscopy and electron tomography (ET) with micro-contact printed circular fibronectin patterns to show that self-organization of the actin cytoskeleton contains a built-in machinery that allows cells to develop left–right asymmetry (2).

ECM protein micropatterning has also been used together with traction force microscopy (TFM) as a tool in mechanobiology research. Oakes et al. confined single cells to ECM protein islands on gels and showed that total strain energy increased with cell area, irrespective of cell shape, number of focal adhesions, and substrate stiffness (9). Moreover, local curvature of the cell, dictated by the ECM protein micropattern, regulated the distribution of traction stress (9). Schaumann et al. found similar behavior in epithelial colonies confined to ECM protein micropatterns (10). Tseng et al. and Sim et al. used TFM together with micropatterning to infer cell-cell forces across cell pairs on ECM micropatterns (11, 12). Spatial organization of the ECM regulated positioning of cell-cell junctions (11) and cell-cell forces increased with increasing aspect ratio for rectangular micropatterns (12). Liu et al. used ECM micropatterning of microfabricated traction force sensors to correlate cell-cell forces with adherens junction size (13). While micropatterning together with TFM helps to relate spatial constraints to cell-cell and cell-substrate mechanics, the nanometer-scale protein organization underlying mechanotransduction, changes in morphology, and cell mechanics has not been observed.

Cryogenic electron tomography (cryo-ET) enables structural characterization of macromolecular complexes in intact vitrified cells in their native state (14–19). Correlative light and electron microscopy (CLEM) technologies can directly tie high-resolution structural information from cryo-ET to spatio-temporal dynamics of cellular events at spatially defined regions of the cell (20–23). A prerequisite to using CLEM technology for the study of mechanotransduction is the availability of cell culture substrates amenable to cryo-ET that can also constrain the morphology of the cells to be imaged.

Manual micropatterning via micro-contact printing of ECM proteins on cell-culture substrates and on transmission electron microscopy amenable substrates (EM grids) has been used successfully to apply CLEM and cellular tomography in the context of spatially confined cell spreading (2). In micro-contact printing, a molecular ink (i.e., protein) is placed onto a substrate using a soft polymer “stamp”, typically cast from a photolithographically-defined topographical mold on a silicon wafer (3, 24–28). Although the process is straightforward once the master mold is made, efficient and high-resolution patterning subsequently depends on achieving conformal contact between the deformable stamp and the substrate (25), and clean detachment of the stamp from the substrate following contact (29). Protein transfer is frequently aided by the application of force to the stamp (30, 31), which may damage fragile substrates. Micro-contact printing has been well characterized for protein transfer to relatively rigid, continuous substrates such as glass coverslips (32, 33). However, a systematic study of protein transfer to EM grids has not been performed.

Here, we describe an alternative robust and reproducible method for depositing ECM protein micropatterns on EM grids, which eliminates the need for physical contact with the EM grid surface to be patterned. For maskless photopatterning of EM grids, we employed the Alvéole PRIMO maskless UV-patterning system (Alvéole, Paris, France) that enables multi-protein photopatterning by light-induced molecular adsorption (34). With this technique, we produced large areas of uniform ECM micropatterns that enabled attachment of various mammalian cells to specific locations on EM grids. By controlling the pattern size and seeding density, the technique can be optimized to yield dominant instances of single cells or cell doublets to control the size and position of cell-cell contacts, as shown by Tseng et al. (11). In addition, the use of virtual masks makes the tool suitable for rapid prototyping, for example to accommodate different pattern areas matched to cell type differences in overall volume during cell spreading. For studies where precise aspect ratios or use of EM grids are required, maskless photopatterning can serve as a more reliable patterning technique to aid mechanobiology studies relating cell morphology and cell mechanics.

## MATERIALS AND METHODS

### Maskless photopatterning

We performed maskless photopatterning on EM grids amenable for mammalian cell culture and cryo-ET (Fig. 1a). EM grids consist of a thin gold mesh, 3.05 mm in diameter, overlaid with a perforated (“holey”) or continuous hydrophobic carbon thin film (35–38). In this study we used 200 mesh gold grids overlaid with 12 nm thick carbon film (Quantifoil Holey Carbon Film Q225AR-520, Quantifoil Micro Tools GmbH, Großlöbichau, Germany), featuring 5 μm wide holes arranged orthogonally with a period of 25 μm.

**Figure 1:**
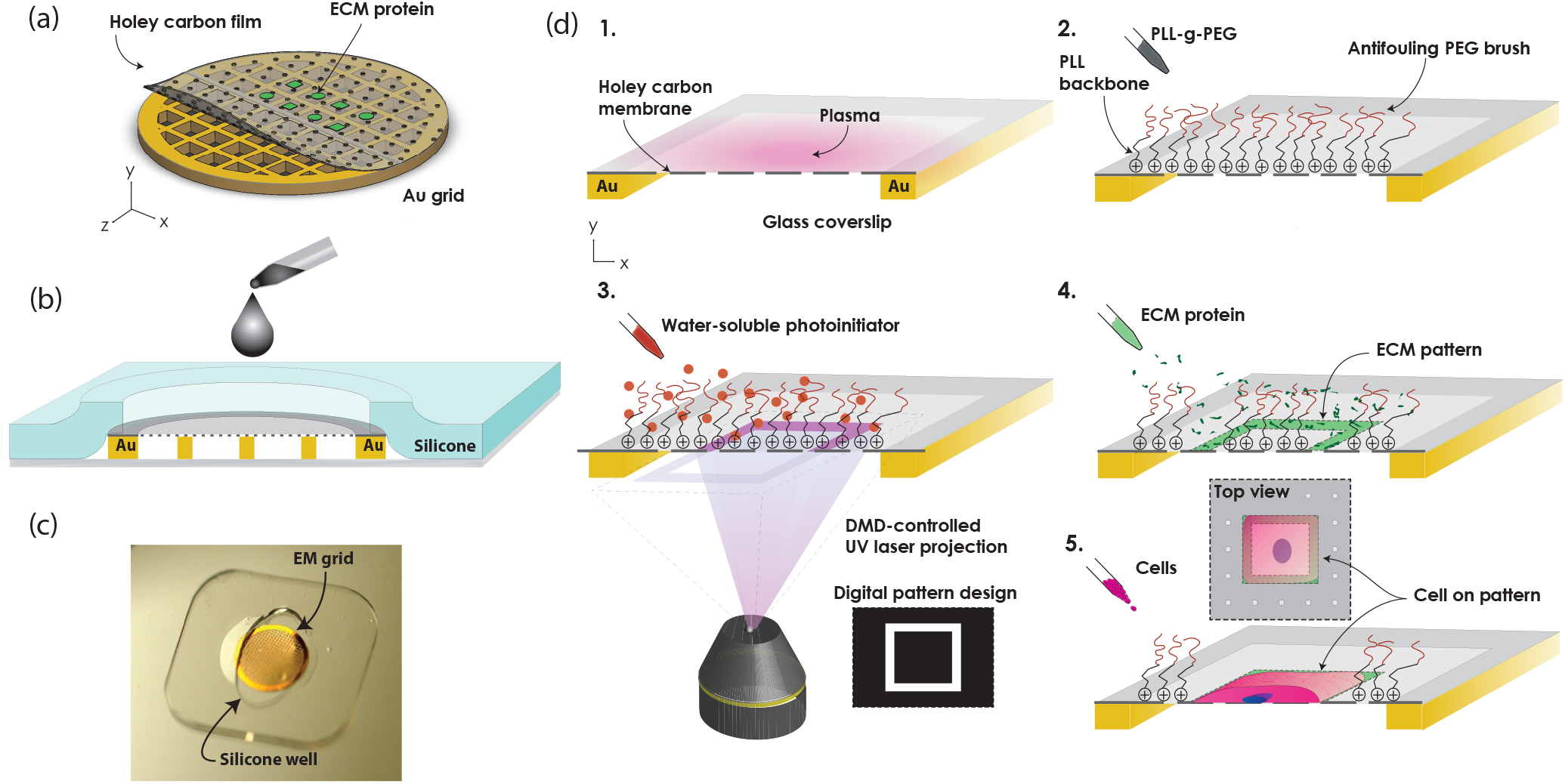
Maskless photopatterning of EM grids. (a) Schematic shows ECM micropatterns aligned to suspended windows of holey carbon thin film on an EM grid. (b) Schematic cross-section of silicone stencil assembled with grid on a coverslip shows how it constrains the grid and acts as a well. (c) Photograph of gold EM grid on a glass coverslip, under a silicone stencil. (d) Process flow for EM grid maskless photopatterning. 1. Expose grid surface to atmospheric plasma. 2. Passivate surface with anti-fouling PEG-brush. 3. Add UV-sensitive photo initiator (PLPP) to the surface. 4. Use Alveole PRIMO system to project the digital maskless micropattern with UV light through holey carbon on EM grid, resulting in the local photodegradation of the anti-fouling PEG-brush. 5. Rinse with PBS and incubate with protein, which selectively adsorbs onto the deprotected are of the carbon film. 6. Seed cells (not to scale).

To prepare holey carbon EM grids for maskless photopatterning, we first immobilized the grids carbon-side up on clean glass coverslips using custom silicone stencils (Fig. 1b,c). The stencil constrained the edges of the grid while leaving the surface exposed for the chemical functionalization and photopatterning steps. We tried a variety of different shaped stencils (i.e. circles, ovals, squares, and crosses) before choosing ovals. The ovals were convenient because they held the grids securely in place at two ends, while allowing liquids to be pipetted into the well at the gaps, instead of directly onto the carbon film surface of the EM grid. The oval-shaped wells were cut from a prefabricated 127 or 250 μm thick silicone sheets (Specialty Manufacturing, Inc., Saginaw, MI, USA) using a Silhouette CAMEO 3 electronic desktop cutting machine (Silhouette America, Inc., UT, USA).

The process for ECM micropatterning of EM grids is summarized in Fig. 1d. We mounted the grids on glass coverslips using a silicone stencil and exposed the grid surface to 10 sec of atmospheric plasma at 25-30 W (PE-50, Plasma Etch Inc., Carson City, NV, USA) to render the carbon film surface hydrophilic. We then incubated the grids i n 1 00 μg/ml poly(l-lysine)-graft-poly(ethylene glycol)(PLL(20)-g[3.5]-PEG(2))(SuSoS AG, Dübendorf, Switzerland) in phosphate buffered saline (PBS) for 1 hr at room temperature to electrostatically adsorb PLL-g-PEG polymeric brushes to the surface via the positively charged poly(l-lysine) backbone. The densely packed PEG side chains provide the grid surface with anti-fouling properties against protein adsorption, i.e., biopassivation. We rinsed each grid three times in PBS, then deposited 7-10 μL of a UV-sensitive photoinitiator (PLPP, Alvéole, Paris, France) onto the grid. We then placed the coverslips or glass-bottom dishes holding the grids on the stage of a Leica DMi8 outfitted with an Alvéole PRIMO maskless UV-patterning system.

We used open source graphics software, Inkscape (39), and ImageJ (40) to generate binary 8-bit mask image files that we loaded into PRIMO’s control software, Leonardo. The software displays a preview of the mask image to be patterned onto a live image of the substrate surface obtained from the microscope camera, allowing for precise alignment of the patterns onto the suspended portion of the carbon film of the EM grid, including with the micron-sized holes of the holey carbon. Once alignment of the pattern to the grid was complete, the pattern was projected through the carbon film by the PRIMO system, using a 5.2 mW, 375 nm UV laser and a digital micromirror array. The projected UV-light pattern results in the localized photodegradation of the antifouling PLL-g-PEG brush. A dose of 2500 mJ/mm^2^ was sufficient to obtain complete photodegradation of the PEG brush on the carbon film surface of the EM grid. Using this process, 36 replicas of a pattern containing four distinct shapes (Fig. 2) could be achieved in 40 min. Following UV exposure, we rinsed the grids three times in PBS and incubated them for 1 hr at room temperature with a fluorescently labeled ECM protein solution at a dilution of 100 μg/ml in PBS to adsorb the protein to the photo-deprotected regions of the grid. The ECM proteins used in this study included 488 Oregon green gelatin (Fig. 2a; G13186, Thermo Scientific, Waltham, MA, USA) and rhodamine labeled fibronectin (Fig. 2b; FNR01, Cytoskeleton, Denver, CO, USA). Excess protein was rinsed off with PBS prior to cell seeding. We kept the grids hydrated at all times throughout the duration of this procedure so as not to compromise the integrity of the grid support film as well as the biopassivating properties of the PLL-g-PEG.

**Figure 2:**
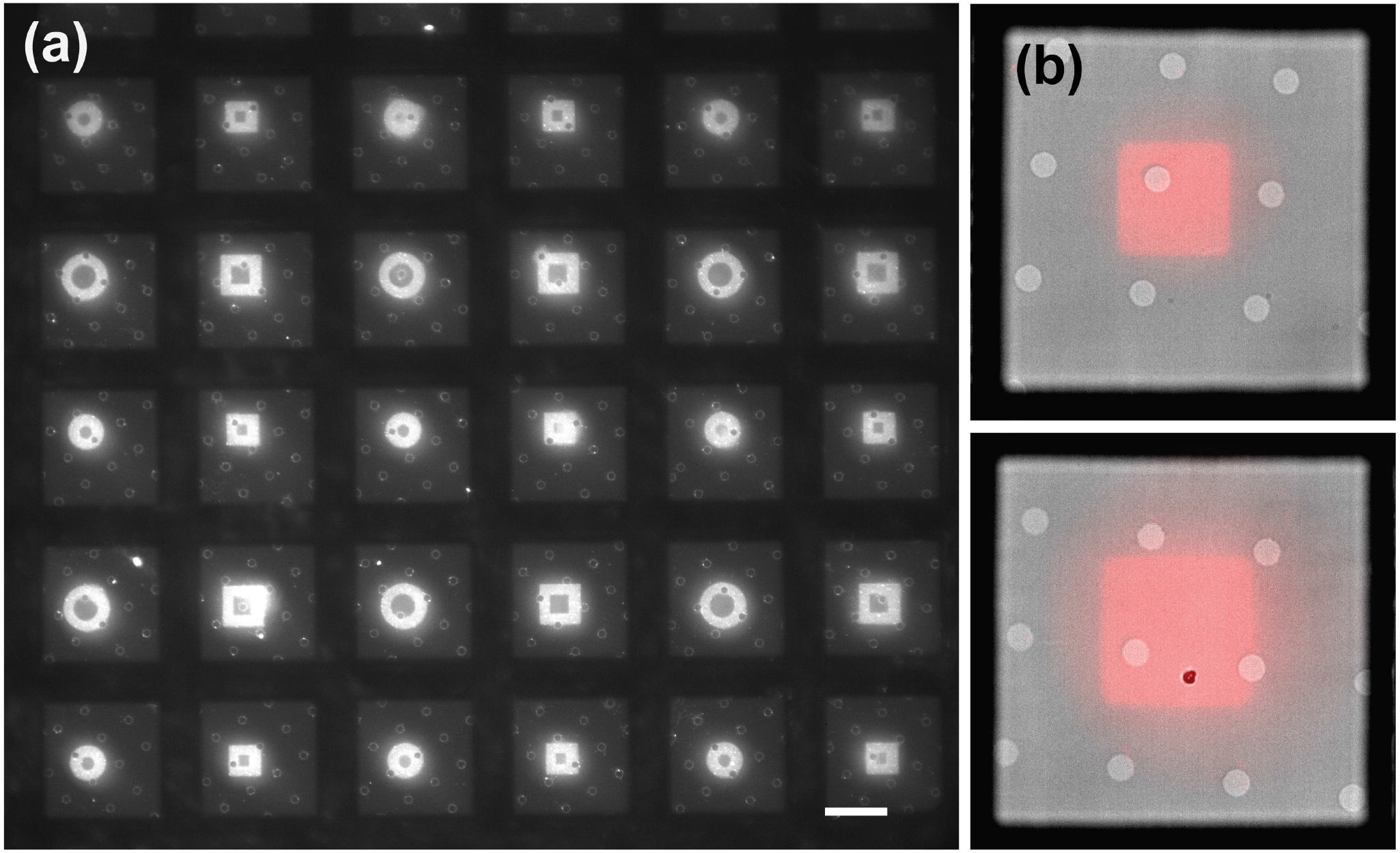
High resolution, large area ECM protein micropatterns of arbitrary geometry patterned on holey carbon EM grids with maskless photopatterning. Squares have side lengths of 24.8 and 32.4 μm. Circles are 27.9 and 36.6 μm in diameter. The shapes are positioned between the gold grid bars of 200 mesh EM grids. (a) Hollow Oregon green 488-gelatin squares and circles. Line width is 8 μm. Scale bar, 50 μm. (b) Overlay of brightfield and fluorescent micrographs show continuous rhodamine-fibronectin square patterns on grids with holey carbon. Holes in carbon 5 μm in diameter, with 20 μm spacing.

To assess the potential damage to the carbon film substrate caused by UV photopatterning, we used an inverted Nikon Ti-E microscope with a 20x Plan Apo Lambda air objective lens to count the number of grid squares (mesh) covered by intact holey carbon film before and after micropatterning. We evaluated four central regions comprised of 6×6 grid squares for each of the EM grids.

In addition to the suspended holey carbon film of EM grids, we also produced ECM micropatterns on continuous carbon thin films deposited on glass coverslips, following the same general procedure described above. However, we modified the substrate configuration so that the carbon-coated coverslip was face down above another glass coverslip with a silicone well containing PLPP acting as a spacer between them.

### Micro-contact printing

We compared the quality of the micro-contact printed protein patterns to protein patterns generated via maskless photopatterning by patterning 488 Oregon green gelatin rectangles using both methods on carbon coated coverslips. Polydimethylsiloxane (PDMS) stamps (Sylgard 184 PDMS, Dow, Midland, MI, USA) were cast from SU-8 photoresist molds on a silicon wafer. The SU-8 was patterned on the wafers using standard photolithography through transparency masks (CAD/ART Services, Inc., Bandon, OR, USA) as described elsewhere (24). We mounted the PDMS stamps on glass coverslips and incubated them with gelatin at a concentration of 100 μg/ml in PBS at room temperature for 1 hr. We then aspirated the excess protein, rinsed the stamps in PBS, and dried them under a stream of nitrogen gas. We brought the stamps into contact with the carbon surfaces and added 50 g weights to the back of the stamps. After 2 min, we removed the weights and left the stamps in contact with the carbon-coated coverslips for another 3 min. Following this, we removed the stamps and placed the samples in PBS for imaging. Following micro-contact printing, the carbon coated coverslips required incubation in PLL-g-PEG at a concentration of 100 μg/ml for 1 hr (biopassivation) to prevent cell spreading outside the area of the ECM patterns. We also micro-contact printed gelatin on EM grids with two modifications of the above protocol. First, to prevent grid damage, we did not apply force to the backs of the stamps using weights. When this did not result in sufficient protein transfer, we included the weights and left out the protein rinsing step prior to making contact between the stamp and the grid. Following micro-contact printing, EM grids were screened for physical integrity and pattern quality.

### Analysis of protein patterns

We acquired images of rectangular protein patterns on carbon-coated coverslips generated by maskless photopatterning and micro-contact printing at 40x magnification and processed them using ImageJ (40). We created a stack by using the cvMatch_Template plugin (41), and two macros (https://github.com/adenisin/ImageJMacros) (30). We aligned the images in the stack using the Align slices in stack plugin (41) and binarized each image in the stack using Otsu thresholding. We averaged the images in the stack and fit circles to the corners of the resulting images to measure radius of curvature.

### Cell culture

We used a variety of mammalian cell types to test cell viability, spreading and spatial confinement on maskless photopatterned EM grids. Madin-Darby Canine Kidney strain II (MDCKII) cells were transfected with Life-Act GFP using the Amaxa Biosystem Nucleofector II system (Lonza, VCA-1005) (Gift from Jens Moller, ETH Zurich, and Teemu Ihalainen, BioMediTech). We cultured these cells in low glucose DMEM (ThermoFisher 11885084) supplemented with 10% FBS, 1% penicillin-streptomycin (PenStrep, ThermoFisher 15140122), and 0.25 mg/ml G418 selection antibiotic (Sigma-Aldrich, G418-RO Roche). Parental Potorous tridactylus Kidney 1 (PtK1) cells (ATCC CRL-6493) were cultured in the same type of media used for MDCKII cells, without selection antibiotic.

Human foreskin fibroblast cells (HFF)(CCD-1070Sk ATCC CRL-2091, American Type Culture Collection, Manassas, VA, USA) were cultured in high glucose DMEM without phenol red (21063-045, Life technologies, Carlsbad, CA), supplemented with 10% FBS, 1x MEM Non-essential amino acids (11140050, Gibco), 1x sodium pyruvate (Corning, MT-25-000-CI), and 1% penicillin-streptomycin.

Our initial troubleshooting assays were confined to micropatterned surfaces generated on continuous carbon film on glass slides. These served to establish (1) the viability of both epithelial cell types on the patterned substrates (Fig. S1 in the Supporting Material) and (2) the necessity of plasma treating carbon surfaces for efficient biopassivation of carbon surfaces with PLL-g-PEG (Fig. S2). In preparation for cell seeding on EM grids patterned with ECM proteins, MDCKII or PtK1 cells from cultures in logarithmic phase were trypsinized and resuspended in culture medium containing 10% serum to inactivate trypsin. Cells were seeded onto EM grids and allowed to adhere for 12-16 hr, prior to fixation and imaging as described previously (22, 42). HFF cells were plated on patterned grids, incubated for 1 hr, imaged live for approximately 1 hr and fixed after another incubation period of 0.5 hr. They were fixed by incubation in a solution of 4% paraformaldehyde (AA433689M, Alfa Aesar, Haverhill, MA, USA) in PBS for 15 min at room temperature.

### Imaging

Epifluorescence imaging of protein patterns, HFF, and MDCKII cells was performed on an inverted Nikon Ti-E microscope (Nikon, Minato, Tokyo, Japan) equipped with a Heliophor light engine (89 North) and an Andor sCMOS Neo camera using a 20x Plan Apo Lambda air objective lens, a 40x Plan Apo Lambda air objective lens and a 60x Plan Apo Lambda oil objective lens. Live cell imaging was performed at 37 °C in cell culture media. Fixed cells were imaged in PBS.

Additional epifluorescence images of protein patterns, and PtK1 cells were acquired on an inverted light microscope (Eclipse TE 2000-U, Nikon) equipped with a manual controlled shutter, filter wheels, and a 14-bit cooled CCD camera (Orca II) controlled by MetaMorph software (Universal Imaging) by a Plan Fluor ELWD 40/0.60 Ph2 or Plan Fluor 10/0.30 Ph1 objective lens (Nikon).

Confocal images of LifeAct GFP MDCKII cells on carbon coated coverslips were acquired on a Leica SP8 confocal microscope (Leica Microsystems Inc., Buffalo Grove, IL, USA). Z-stack slices ranged from 0.15 to 0.20 μm and were processed using Leica’s Lightning deconvolution package.

When imaging maskless photopatterned EM grids that are mounted on the glass coverslip used as support surface for patterning, the user will observe an apparent “double image” of the same pattern at two z heights (representing the planes of the glass and the suspended holey carbon thin film). For fluorescent microscopy of patterned grids, it is helpful to identify the holes in the holey carbon film to differentiate the correct focal plane of the grid from that of the glass coverslip below (Fig. S3).

## RESULTS AND DISCUSSION

Here, we describe a robust and reproducible method for depositing ECM protein micropatterns on EM grids. Our goal in developing this process is to ultimately enable use of CLEM and cryo-ET to correlate changes in nanometer-scale organization at cell-cell and cell-ECM contacts with strain energy states and traction stress distribution in the cell.

We found that the Alvéole PRIMO maskless UV-patterning system is amenable to patterning EM grids using the setup detailed in the Methods section. We reproducibly micropatterned large areas of EM grids with uniform protein patterns of different shapes and sizes (Fig. 2 and Fig. S4) and were able to reproducibly position micropatterns on the grids (Fig.2). In addition, maskless photopatterning did not substantially compromise grid integrity. An average of only 0.3+/-0.2 (S.E.M.) grid squares per each assessed 36 grid-square region (less than 1%) were damaged following maskless photopatterning (n=12, Table S1, Fig. S5).

### Comparison of maskless photopatterning to other ECM micropatterning techniques

We compared maskless photopatterning to photoresist lift-off-assisted micropatterning (30) on EM grids. In photoresist lift-off-assisted micropatterning, the substrate to be patterned (i.e., glass slide) is coated with photoresist, which is then patterned by photolithography, and incubated in PLL-g-PEG to create an anti-fouling coating on the surface (30). When the patterned photoresist is removed in organic solvent, the PLL-g-PEG on the surface remains intact and the newly bare surface can be backfilled with protein. We were able to successfully align photoresist patterns to EM grids (Fig. S6), but found the EM grids incompatible with the organic solvent (N-Methyl-2-Pyrrolidone) used during the lift-off stage. Modification of this technique to employ a more compatible organic solvent could be pursued in the future.

We next compared the resolution of the maskless photopatterning method to micro-contact printing on carbon, by patterning rectangles on continuous carbon thin films mounted on glass coverslips using both methods. While micro-contact printing is a simpler, more accessible, and lower-cost patterning method than maskless photopatterning, maskless photopatterning of continuous carbon films gave a higher yield of continuous micropatterns, better protein uniformity, and higher patterning resolution. There were entire regions of the micro-contact-printed carbon coated coverslips that contained no ECM micropatterns or non-continuous micropatterns after contact with an ECM-laden stamp (Fig. S7). In addition, our results suggest that for studies where precise aspect ratio is required, maskless photopatterning might be a more reliable method.

Fig. 3 shows the averages of binarized protein patterns on continuous carbon films generated via maskless photopatterning (n=70) and micro-contact printing (n=112) of fluorescent gelatin as compared to a rectangular template featuring a 1:7 aspect ratio rectangle with a 1500 μm^2^ area. The corners of the rectangles generated by maskless photopatterning were sharper than the micro-contact printed patterns. Using ImageJ, we measured an average radius of 1.4 μm at the corners of maskless photopatterned rectangles and an average radius of curvature of more than double that, 3.8 μm, for the micro-contact printed rectangles. Upon examination, the features on the PDMS stamps had rounded edges (Fig. S8), leading us to believe that the lower resolution was caused by aberration of ultraviolet light during photolithography or release of the PDMS stamp from the master (32). The resolution of micro-contact printing on carbon films can be improved by using a higher quality photomask for photolithography, but this would drive up the cost per design iteration. In addition, our results are consistent with a previous study which compared micro-contact printing and photoresist lift-off-assisted micropatterning of ECM micropatterns transferred to gels that were generated using photomasks of equal resolution and quality (30). The micropatterns transferred from glass coverslips that underwent photolithography had higher fidelity to the shape of the template than micropatterns transferred from the micro-contact printed glass coverslips (30).

**Figure 3:**
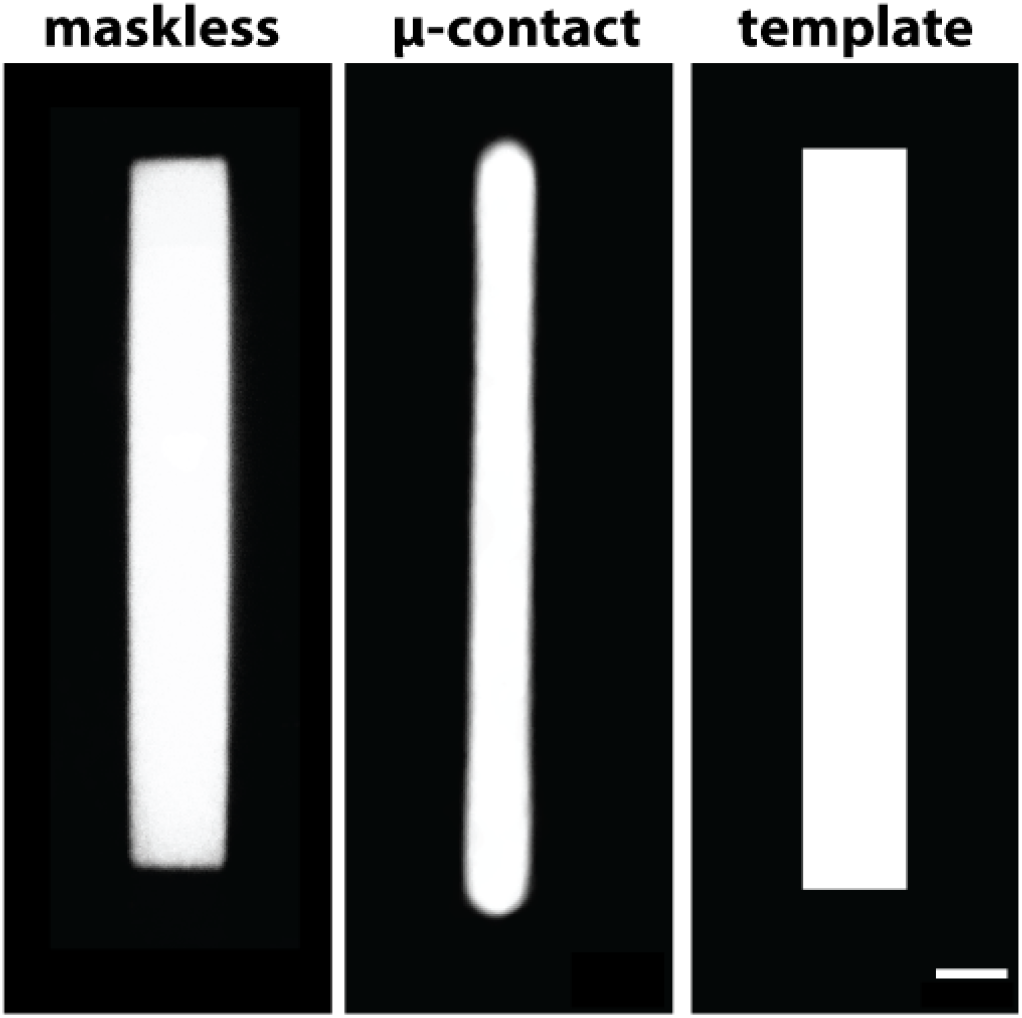
Maskless photopatterning is a more accurate ECM protein micropatterning technique than micro-contact printing. Averaged binarized fluorescent images of Oregon green 488-gelatin rectangles on continuous carbon films produced using maskless photopatterning (n=70) and micro-contact printing (n=112). Right panel shows template. Scale bar, 10μm.

Geometric accuracy of micropatterns, as characterized by sharpness of vertices, is an important parameter for mechanobiology studies and for simulating the spatial confinement a cell might experience in an *in vivo* or *in vitro* environment due to constraints from neighboring cells (43). Kollimada et al. found that epithelial cell confinement to vertices of shapes before removal of the confinement influenced leading edge speeds of cell monolayers (44). Local curvature also regulates the distribution of traction stress (9). Both Rape et al. and Oakes et al. observed higher traction stresses at corners in cells plated on ECM squares (9, 45). Moeller et al. compared MDCKII cell pair behavior on square patterns with sharp and rounded corners, observing increased lamellipodial length and exploratory behavior in MDCKII cell pairs plated on squares with sharper vertices (30). Control over ECM protein pattern aspect ratio is another important parameter for mechanobiology studies. For example, Ribeiro et al. showed that cardiomyocytes differentiated from human pluripotent stem cells (hPSC-CMs) on rectangular matrigel micropatterns of the same area had increased sarcomere activity and myofibril alignment on micropatterns with an aspect ratio of 1:7 (46).

The aspect ratio of the averaged maskless photopatterned rectangles on continuous carbon film was 1:7.3, closely replicating the aspect ratio of the template. The aspect ratio of the micro-contact printed rectangles was 1:14.4, nearly twice that of the template. While the average length of the micro-contact printed patterns was close to the length of the template (3% longer), the aspect ratio was higher due to the smaller width of these micropatterns, which was approximately 50% of the width of the template. The aspect ratio of the features on the stamps used for micro-contact printing more closely resembled the aspect ratio of the template than the ECM micropatterns (Fig. S8), suggesting that further process optimization would be required to achieve the desired micropattern dimensions. For example, a possible strategy could be to design a photomask with a rectangle width that is double the desired ECM pattern width.

The success of manual micro-contact printing of EM grids likely depends on the operator, the grid type, the micropattern shape and density, and the pressure applied by the stamp onto the carbon film. For example, in our experience the carbon film was damaged on average in 20.8+/−2.2 (S.E.M.) squares out of each 36 grid square region following removal of the stamp (n=20, Table S2). In contrast, the average damage to carbon film as a result of maskless photopatterning was significantly lower; 0.3+/−0.2 (S.E.M.) grid squares per each assessed 36 grid-square region (p-value=4.18 × 10^−8^). Maskless photopatterning does not require direct contact with the EM grid surface because it is lithographically generated. Protein patterns that were transferred to EM grids by micro-contact printing were deposited mainly on carbon film above the gold grid bars and not on the suspended carbon mesh (Fig. S9), and we could not align the patterns to the grid using this micropatterning method. In the maskless photopatterning described here, pattern projection from an image file onto an EM grid in real time allows for precise alignment of micropatterns to specific regions of interest on the EM grid, thus maximizing experimental setup through accurate spatial deposition of the patterns, and avoiding grid regions that are not amenable for analysis due to lack of transparency to the electron beam (i.e., grid metal bars; Fig. S9). Finally, rapid prototyping of micropattern design can be performed at negligible cost per design iteration.

### Cells on maskless photopatterned EM grids

We demonstrated that maskless photopatterning can produce viable substrates that promote cell growth and can confine cell spreading to within the boundary of an ECM micropattern. We were able to confine adherent PtK1 (Fig. 4), MDCKII and HFF cells (Fig S10) to pre-deposited ECM islands of different shapes and sizes on EM grids. The maskless photopatterned EM grid in Fig. 4 features hollow squares that are 24.8 and 32.4 μm wide and hollow circles that are 27.9 and 36.6 μm wide, all with a line width of 8 μm. Cells spread on these micropatterns for 12 hr before imaging.

**Figure 4:**
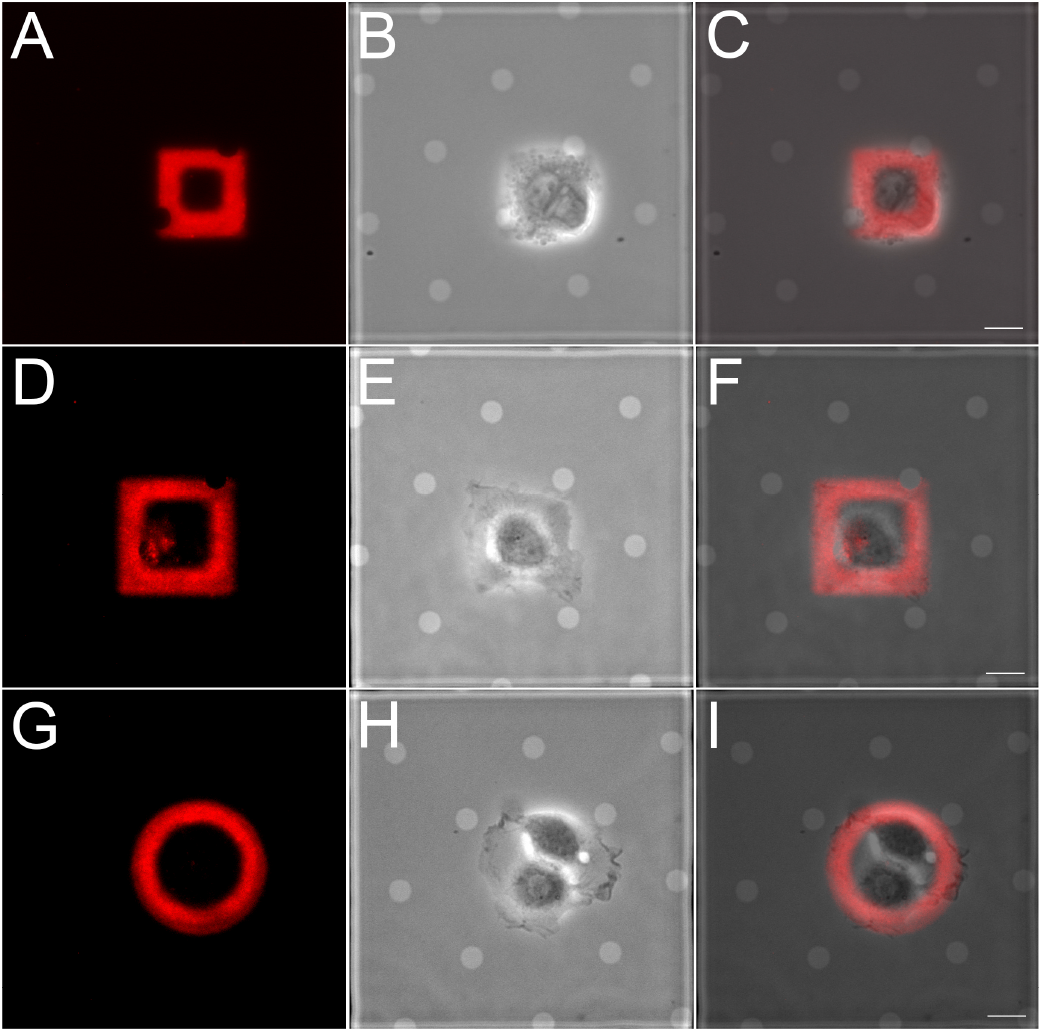
Epithelial cells are confined to ECM patterns on EM grids generated by maskless photopatterning. PtK1 cells were plated on rhodamine-fibronectin square (A-C, 24.8 μm wide; D-F, 32.4 μm wide) and circle (36.6 μm wide) patterns, allowed to adhere and spread for 12 hr. (A, D, G) Rhodamine-fibronectin; (B, E, H) phase contrast; (C, F, I) overlay. Scale bar, 10μm.

Cells were consistently confined to ECM micropatterns generated by maskless photopatterning on carbon thin films, although we observed that they did not always spread across the entire area of the micropattern. For example, although MDCKII cells have spread fully on ECM shapes of comparable area (Fig. S11, S12) (12), the cell doublet pictured on the fluorescent gelatin circle in Fig. S10a did not fill the entire micropatterned circle, even after 16 hr of incubation. We qualitatively observed that patterning the outline of a shape (i.e., 8 μm wide lines forming a 35 × 60 μm rectangle as in Fig. S13) vs. a filled shape encouraged cell spreading across the ECM shape by limiting spreading to peripheral regions of the shape where ECM is present. HFF cells spread more quickly than the two epithelial cell lines and spread across such ECM micropatterns after only 2.5 hr incubation (see Supporting Movies).

By prescribing cell morphology on EM grids, maskless photopatterning can enable mechanobiology studies that correlate gross morphological changes with spatially localized changes in ultrastructure organization at the nanometer scale. This approach, facilitated by the ease of changing the pattern’s physical dimensions via the Alveole PRIMO maskless UV-patterning system, could, for example, serve as basis for mechanobiology studies of cell-cell junctions. Several recent papers quantified traction stresses in cells plated on E-cadherin coated substrates of varying stiffness to investigate cadherin based rigidity sensing at cell-cell contacts (47, 48). The maskless platform can provide a basis for further probing the protein structures underlying mechanotransduction at cell-cell contacts. Maskless photopatterning, using the sample configuration presented here, could facilitate and increase the success rate of cryo-ET data acquisition through positioning specific areas of interest of cells a favorable imaging positions. We expect maskless photopatterning to be compatible with cryogenic focused ion beam (cryo-FIB) thinning of vitrified mammalian cells (49) and suggest that fluorescent protein micropatterns may be used to guide and select regions of a vitrified sample for cryo-FIB.

## CONCLUSION

Here we demonstrated a robust, versatile, and non-destructive maskless photopatterning technology for depositing ECM proteins on cryo-EM grids with programmed shape. We showed that cells plated on maskless photopatterned ECM micropatterns on EM grids are viable on and confined to the micropatterns. In addition, the maskless photopatterned micropatterns were uniform, and could be positioned precisely at specific locations on EM grids. Robustness of the pattern transfer and reproducibility of pattern alignment to EM grids increase the chances of achieving successful cryo-tomograms of cells on ECM patterns. Demonstrating these capabilities for maskless photopatterning of EM grids constitutes progress toward using ECM micropatterning in conjunction with cryo-ET. We anticipate maskless photopatterning of EM grids to be compatible with CLEM, vitrification and cryo-FIB, to enable researchers to program cell shape, strain energy, and cell location on EM grids for high-resolution mechanobiology studies.

## Supporting information

HFF cells spreading on Micropatterns

## AUTHOR CONTRIBUTIONS

LE, GG, GP, NV, WIW, DH, and BLP designed research; LE, GG, MFS and LPD performed research; LE, GG, and LPD analyzed data; NV contributed analytical tools for CLEM work. LE, GG, LPD and GP wrote the paper. NV, WIW, DH and BLP edited the paper.

## ACKNOWLEDGMENTS

Thanks to members of the Pruitt, Dunn, Weis, Nelson, Volkmann and Hanein labs for helpful advice and discussion, and to A.K. Denisin, R.E. Wilson, and A.R. Dunn (ARD) in particular. Work was performed in part in the nano@Stanford labs, which are supported by the National Science Foundation (NSF) as part of the National Nanotechnology Coordinated Infrastructure under award ECCS-1542152 (BP). This work was supported by NSF CMMI 1662431 (BLP) and National Institutes of Health R01GM119948 (DH, WW, NV, BLP), a Stanford ChEM-H fellowship (LE) and National Institutes of Health R01HL128779 (AD). Work performed at SBP was in part supported by National Institutes of Health grants S10-OD012372 (DH) and P01-GM098412 (DH) towards purchase of the correlated light and TEM studies. This work used the NRI-MCDB Microscopy Facility and the Resonant Scanning Confocal Microscope supported by NSF MRI grant DBI-1625770.

## REFERENCES

1. Théry, M., 2010. Micropatterning as a tool to decipher cell morphogenesis and functions. J Cell Sci 123:4201–4213.

2. Tee, Y. H., T. Shemesh, V. Thiagarajan, R. F. Hariadi, K. L. Anderson, C. Page, N. Volkmann, D. Hanein, S. Sivaramakrishnan, M. M. Kozlov, et al., 2015. Cellular chirality arising from the self-organization of the actin cytoskeleton. Nature cell biology 17:445.

3. Singhvi, R., A. Kumar, G. P. Lopez, G. N. Stephanopoulos, D. Wang, G. M. Whitesides, and D. E. Ingber, 1994. Engineering cell shape and function. Science 264:696–698.

4. Chen, C. S., M. Mrksich, S. Huang, G. M. Whitesides, and D. E. Ingber, 1997. Geometric control of cell life and death. Science 276:1425–1428.

5. Watt, F. M., P. W. Jordan, and C. H. O’Neill, 1988. Cell shape controls terminal differentiation of human epidermal keratinocytes. Proceedings of the National Academy of Sciences 85:5576–5580.

6. Lee, J., A. A. Abdeen, D. Zhang, and K. A. Kilian, 2013. Directing stem cell fate on hydrogel substrates by controlling cell geometry, matrix mechanics and adhesion ligand composition. Biomaterials 34:8140–8148.

7. Song, W., H. Lu, N. Kawazoe, and G. Chen, 2011. Adipogenic differentiation of individual mesenchymal stem cell on different geometric micropatterns. Langmuir 27:6155–6162.

8. Théry, M., V. Racine, A. Pépin, M. Piel, Y. Chen, J.-B. Sibarita, and M. Bornens, 2005. The extracellular matrix guides the orientation of the cell division axis. Nature cell biology 7:947.

9. Oakes, P. W., S. Banerjee, M. C. Marchetti, and M. L. Gardel, 2014. Geometry regulates traction stresses in adherent cells. Biophysical journal 107:825–833.

10. Schaumann, E. N., M. F. Staddon, M. L. Gardel, and S. Banerjee, 2018. Force localization modes in dynamic epithelial colonies. Molecular biology of the cell 29:2835–2847.

11. Tseng, Q., E. Duchemin-Pelletier, A. Deshiere, M. Balland, H. Guillou, O. Filhol, and M. Théry, 2012. Spatial organization of the extracellular matrix regulates cell–cell junction positioning. Proceedings of the National Academy of Sciences 109:1506–1511.

12. Sim, J. Y., J. Moeller, K. C. Hart, D. Ramallo, V. Vogel, A. R. Dunn, W. J. Nelson, and B. L. Pruitt, 2015. Spatial distribution of cell–cell and cell–ECM adhesions regulates force balance while maintaining E-cadherin molecular tension in cell pairs. Molecular biology of the cell 26:2456–2465.

13. Liu, Z., J. L. Tan, D. M. Cohen, M. T. Yang, N. J. Sniadecki, S. A. Ruiz, C. M. Nelson, and C. S. Chen, 2010. Mechanical tugging force regulates the size of cell–cell junctions. Proceedings of the National Academy of Sciences 107:9944–9949.

14. Asano, S., B. D. Engel, and W. Baumeister, 2016. In situ cryo-electron tomography: a post-reductionist approach to structural biology. Journal of molecular biology 428:332–343.

15. Beck, M., and W. Baumeister, 2016. Cryo-electron tomography: can it reveal the molecular sociology of cells in atomic detail? Trends in cell biology 26:825–837.

16. Gan, L., and G. J. Jensen, 2012. Electron tomography of cells. Quarterly reviews of biophysics 45:27–56.

17. Medalia, O., I. Weber, A. S. Frangakis, D. Nicastro, G. Gerisch, and W. Baumeister, 2002. Macromolecular architecture in eukaryotic cells visualized by cryoelectron tomography. Science 298:1209–1213.

18. Mahamid, J., S. Pfeffer, M. Schaffer, E. Villa, R. Danev, L. K. Cuellar, F. Förster, A. A. Hyman, J. M. Plitzko, and W. Baumeister, 2016. Visualizing the molecular sociology at the HeLa cell nuclear periphery. Science 351:969–972.

19. Baker, L. A., M. Grange, and K. Grünewald, 2017. Electron cryo-tomography captures macromolecular complexes in native environments. Current opinion in structural biology 46:149–156.

20. Sartori, A., R. Gatz, F. Beck, A. Rigort, W. Baumeister, and J. M. Plitzko, 2007. Correlative microscopy: bridging the gap between fluorescence light microscopy and cryo-electron tomography. Journal of structural biology 160:135–145.

21. Schorb, M., L. Gaechter, O. Avinoam, F. Sieckmann, M. Clarke, C. Bebeacua, Y. S. Bykov, A. F.-P. Sonnen, R. Lihl, and J. A. Briggs, 2017. New hardware and workflows for semi-automated correlative cryo-fluorescence and cryo-electron microscopy/tomography. Journal of structural biology 197:83–93.

22. Marston, D. J., K. L. Anderson, M. F. Swift, M. Rougie, C. Page, K. M. Hahn, N. Volkmann, and D. Hanein, 2019. High Rac1 activity is functionally translated into cytosolic structures with unique nanoscale cytoskeletal architecture. Proceedings of the National Academy of Sciences 116:1267–1272.

23. Hampton, C. M., J. D. Strauss, Z. Ke, R. S. Dillard, J. E. Hammonds, E. Alonas, T. M. Desai, M. Marin, R. E. Storms, F. Leon, et al., 2017. Correlated fluorescence microscopy and cryo-electron tomography of virus-infected or transfected mammalian cells. Nature protocols 12:150.

24. Qin, D., Y. Xia, and G. M. Whitesides, 2010. Soft lithography for micro-and nanoscale patterning. Nature protocols 5:491.

25. Xia, Y., and G. M. Whitesides, 1998. Soft lithography. Angewandte Chemie International Edition 37:550–575.

26. Whitesides, G. M., E. Ostuni, S. Takayama, X. Jiang, and D. E. Ingber, 2001. Soft lithography in biology and biochemistry. Annual review of biomedical engineering 3:335–373.

27. Théry, M., A. Pépin, E. Dressaire, Y. Chen, and M. Bornens, 2006. Cell distribution of stress fibres in response to the geometry of the adhesive environment. Cell motility and the cytoskeleton 63:341–355.

28. Jokhun, D. S., Y. Shang, and G. Shivashankar, 2018. Actin dynamics couples extracellular signals to the mobility and molecular stability of telomeres. Biophysical journal 115:1166–1179.

29. Chen, C. S., M. Mrksich, S. Huang, G. M. Whitesides, and D. E. Ingber, 1998. Micropatterned surfaces for control of cell shape, position, and function. Biotechnology progress 14:356–363.

30. Moeller, J., A. K. Denisin, J. Y. Sim, R. E. Wilson, A. J. Ribeiro, and B. L. Pruitt, 2018. Controlling cell shape on hydrogels using lift-off protein patterning. PloS one 13:e0189901.

31. Shen, K., J. Qi, and L. C. Kam, 2008. Microcontact printing of proteins for cell biology. Journal of visualized experiments: JoVE.

32. Hui, C., A. Jagota, Y. Lin, and E. Kramer, 2002. Constraints on microcontact printing imposed by stamp deformation. Langmuir 18:1394–1407.

33. Bernard, A., E. Delamarche, H. Schmid, B. Michel, H. R. Bosshard, and H. Biebuyck, 1998. Printing patterns of proteins. Langmuir 14:2225–2229.

34. Strale, P.-O., A. Azioune, G. Bugnicourt, Y. Lecomte, M. Chahid, and V. Studer, 2016. Multiprotein Printing by Light-Induced Molecular Adsorption. Advanced Materials 28:2024–2029.

35. Azubel, M., S. D. Carter, J. Weiszmann, J. Zhang, G. J. Jensen, Y. Li, and R. D. Kornberg, 2019. FGF21 trafficking in intact human cells revealed by cryo-electron tomography with gold nanoparticles. eLife 8:e43146.

36. Jasnin, M., S. Asano, E. Gouin, R. Hegerl, J. M. Plitzko, E. Villa, P. Cossart, and W. Baumeister, 2013. Three-dimensional architecture of actin filaments in Listeria monocytogenes comet tails. Proceedings of the National Academy of Sciences 110:20521–20526.

37. Patla, I., T. Volberg, N. Elad, V. Hirschfeld-Warneken, C. Grashoff, R. Fässler, J. P. Spatz, B. Geiger, and O. Medalia, 2010. Dissecting the molecular architecture of integrin adhesion sites by cryo-electron tomography. Nature cell biology 12:909.

38. Gupton, S. L., K. L. Anderson, T. P. Kole, R. S. Fischer, A. Ponti, S. E. Hitchcock-DeGregori, G. Danuser, V. M. Fowler, D. Wirtz, D. Hanein, et al., 2005. Cell migration without a lamellipodium: translation of actin dynamics into cell movement mediated by tropomyosin. The Journal of cell biology 168:619–631.

39. Harrington, B. e. a., 2004–2005. http://www.inkscape.org/.www.inkscape.org.

40. Abràmoff, M. D., P. J. Magalhães, and S. J. Ram, 2004. Image processing with ImageJ. Biophotonics international 11:36–42.

41. Tseng, Q., I. Wang, E. Duchemin-Pelletier, A. Azioune, N. Carpi, J. Gao, O. Filhol, M. Piel, M. Théry, and M. Balland, 2011. A new micropatterning method of soft substrates reveals that different tumorigenic signals can promote or reduce cell contraction levels. Lab on a chip 11:2231–2240.

42. Anderson, K. L., C. Page, M. F. Swift, P. Suraneni, M. E. Janssen, T. D. Pollard, R. Li, N. Volkmann, and D. Hanein, 2017. Nano-scale actin-network characterization of fibroblast cells lacking functional Arp2/3 complex. Journal of structural biology 197:312–321.

43. Fletcher, A. G., M. Osterfield, R. E. Baker, and S. Y. Shvartsman, 2014. Vertex models of epithelial morphogenesis. Biophysical journal 106:2291–2304.

44. Kollimada, S. A., A. H. Kulkarni, A. Ravan, and N. Gundiah, 2016. Advancing edge speeds of epithelial monolayers depend on their initial confining geometry. PloS one 11:e0153471.

45. Rape, A. D., W.-h. Guo, and Y.-l. Wang, 2011. The regulation of traction force in relation to cell shape and focal adhesions. Biomaterials 32:2043–2051.

46. Ribeiro, A. J. S., Y.-S. Ang, J.-D. Fu, R. N. Rivas, T. M. A. Mohamed, G. C. Higgs, D. Srivastava, and B. L. Pruitt, 2015. Contractility of single cardiomyocytes differentiated from pluripotent stem cells depends on physiological shape and substrate stiffness. Proceedings of the National Academy of Sciences 112:12705–12710.

47. Collins, C., A. K. Denisin, B. L. Pruitt, and W. J. Nelson, 2017. Changes in E-cadherin rigidity sensing regulate cell adhesion. Proceedings of the National Academy of Sciences 114:E5835–E5844.

48. Yang, Y., E. Nguyen, G. H. N. S. Narayana, M. Heuze, R.-M. Mege, B. Ladoux, and M. P. Sheetz, 2018. Local Contractions Regulate E-Cadherin Adhesions, Rigidity Sensing and Epithelial Cell Sorting. bioRxiv https://www.biorxiv.org/content/early/2018/12/05/318642.

49. Rigort, A., F. J. Bäuerlein, A. Leis, M. Gruska, C. Hoffmann, T. Laugks, U. Böhm, M. Eibauer, H. Gnaegi, W. Baumeister, et al., 2010. Micromachining tools and correlative approaches for cellular cryo-electron tomography. Journal of structural biology 172:169–179.

