## Supplementary Material for "Extracellular matrix micropatterning technology for whole cell cryogenic electron microscopy studies"

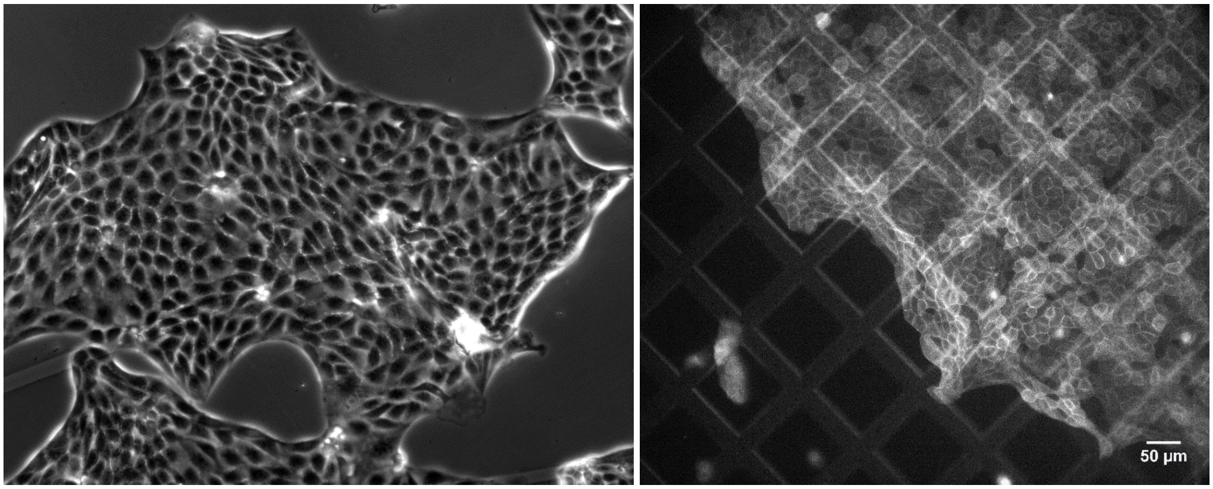

**Fig. S1**: Epithelial cells are viable on continuous carbon films and EM grids. Left: 10x phase micrograph of MDCKII cells on untreated continuous carbon film. Right: Lifeact GFP MDCKII cells plated on EM grids treated with fibronectin, imaged at 10x GFP.

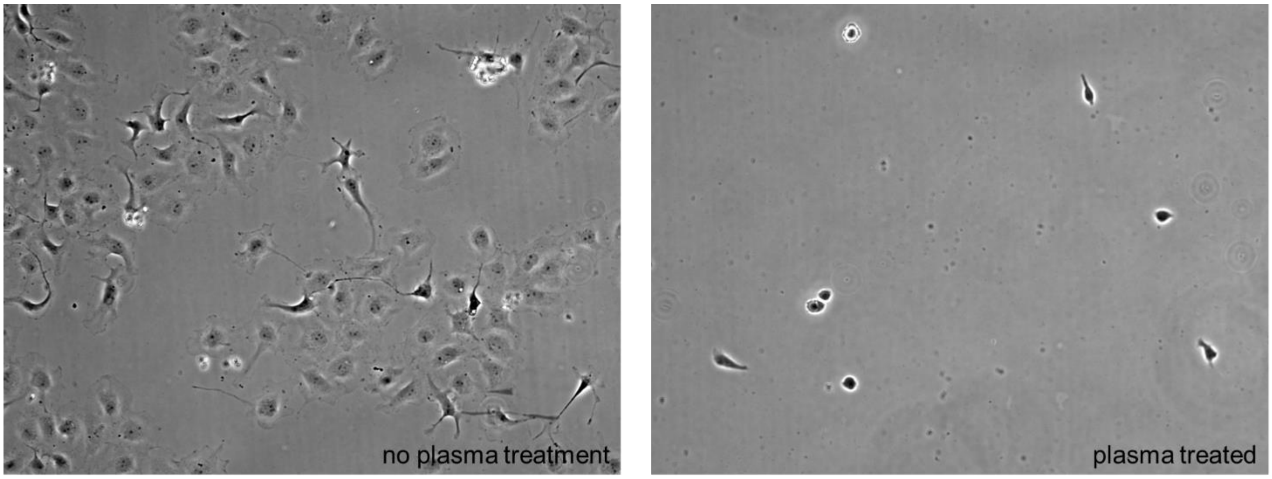

**Fig. S2**: Biopassivation is required to prevent cell adhesion to continuous carbon films. Plasma treatment before PLL-g-PEG incubation (right) is also necessary for biopassivation. Left image shows cells growing on carbon film incubated with PLL-g-PEG, with no prior plasma treatment.

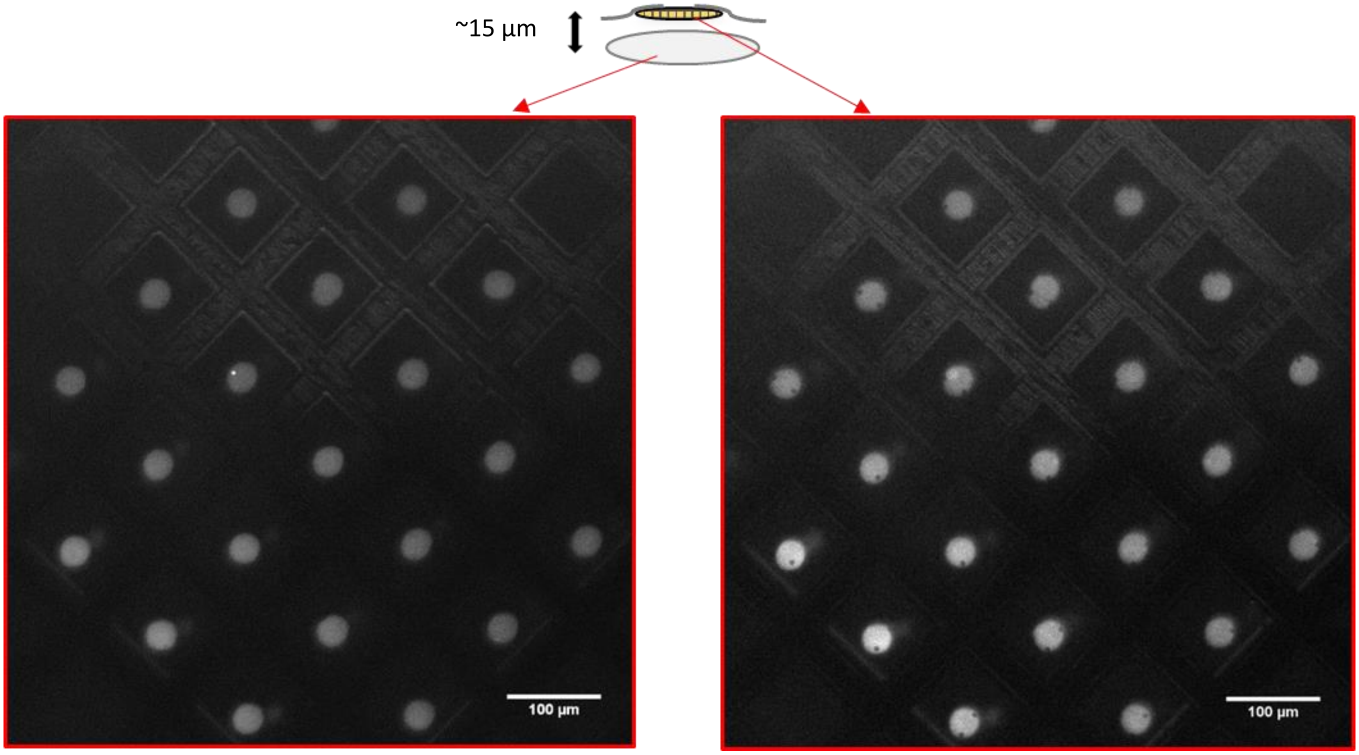

**Fig. S3**: "Double patterning" of Quantifoil Holey Carbon Au EM grid on glass coverslip. The UV laser in the PRIMO system travels through the glass coverslip as well as the carbon film of the EM grids, degrading the PLL-g-PEG on both surfaces. Subsequent protein backfill with ECM then results in two sets of patterns: one on the glass (left) and one on the carbon film (right). This difference can be visualized by the presence of holes in the ECM patterns on the carbon layer. These two surfaces are separated by approximately 15 µm and this phenomenon does not affect the cells' ability to adhere to protein patterns of the carbon surface. The user should be aware of this effect, as not to image the glass layer.

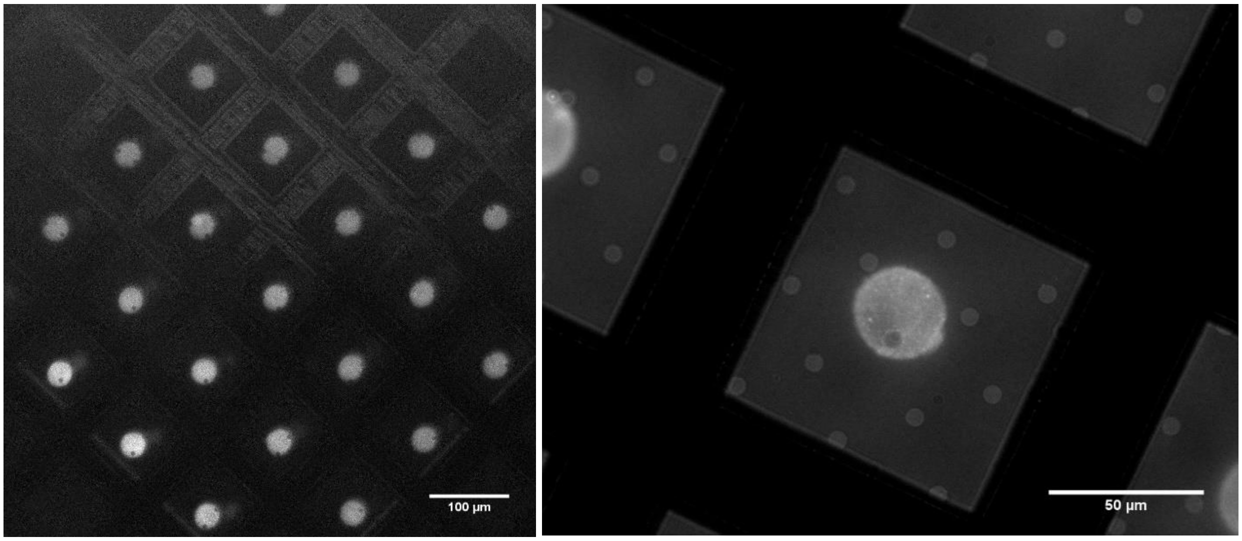

**Fig. S4**: Micropatterned EM grids imaged with fluorescent microscopy. Rhodamine fibronectin islands, 30 µm in diameter (left) were patterned on thin holey carbon Quantifoil Au EM grids. PRIMO can be used to align the patterns within the windows of the metal mesh (left) and consistently demonstrates spatially accurate shapes (right).

| | Broken before | Out of | Broken after |
|---|---|---|---|
| sample1 | 0 | 36 | 0 |
| | 2 | 36 | 2 |
| | 0 | 36 | 0 |
| | 3 | 36 | 4 |
| Sample2 | 16 | 36 | 16 |
| | 5 | 36 | 5 |
| | 0 | 36 | 0 |
| | 7 | 36 | 9 |
| Sample3 | 0 | 36 | 0 |
| | 0 | 36 | 0 |
| | 0 | 36 | 0 |
| | 0 | 36 | 1 |

**Table S1**: Grid windows broken before and after maskless photopatterning.

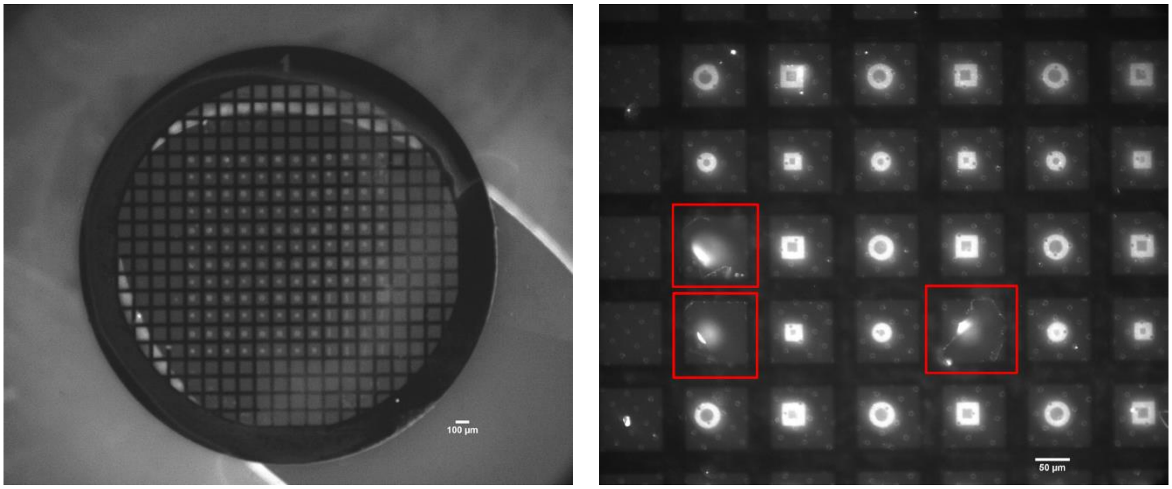

**Fig. S5**: Central 12x12 region of 200 mesh EM grid is patterned with maskless photopatterning. Left: EM grid seen in GFP at 4x magnification. Fluorescent signal is likely from the patterned glass beneath the grid, but features the same pattern as grid. Of the intact grid squares, 1 broke following maskless photopatterning. Of the 144 grid squares, 139 were intact before patterning and 138 had intact holey carbon film after plasma activation and patterning. Oval silicone well can be seen in light gray. Right: EM grid, following maskless photopatterning, imaged in GFP 20x showing three broken grid squares outlined in red (possibly broken before micropatterning).

| | intact before | intact after |
|---|---|---|
| | 35 | 18 |
| | 36 | 16 |
| | 36 | 13 |
| | 36 | 23 |
| | 35 | 6 |
| | 33 | 8 |
| | 33 | 3 |
| | 36 | 6 |
| | 32 | 30 |
| | 36 | 34 |
| | 36 | 31 |
| | 35 | 23 |
| | 36 | 10 |
| | 36 | 2 |
| | 31 | 5 |
| | 36 | 9 |
| | 36 | 10 |
| | 33 | 8 |
| | 32 | 17 |
| | 36 | 8 |
| SUM | 695 | 280 |

**Table S2**: Grid integrity following micro-contact printing. 6x6 Grids square regions observed around central grid region of 200 mesh grid before and after micro-contact printing.

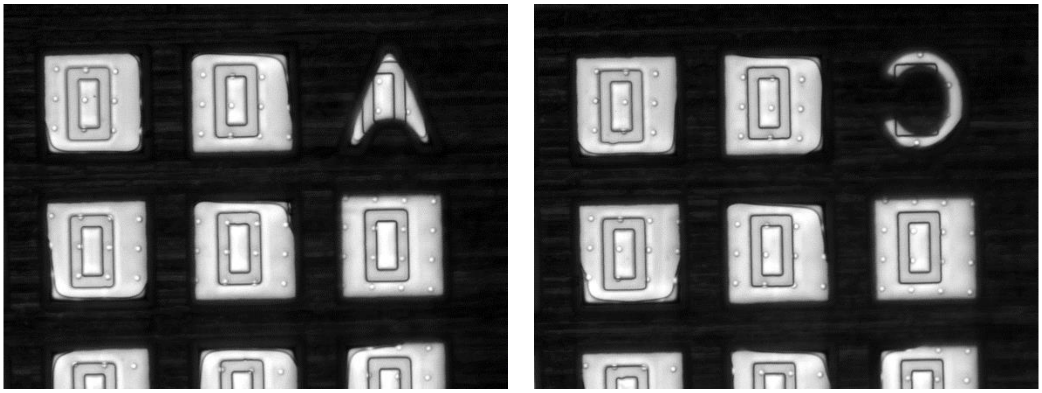

**Fig. S6**: Results of first step of photoresist lift-off assisted micropatterning of EM grids. Alignment of 35x60 micron hollow rectangular photoresist (S1818) patterns to centers of gold finder grids. No holey carbon was present following rinsing in N-Methyl-2-Pyrrolidone (NMP).

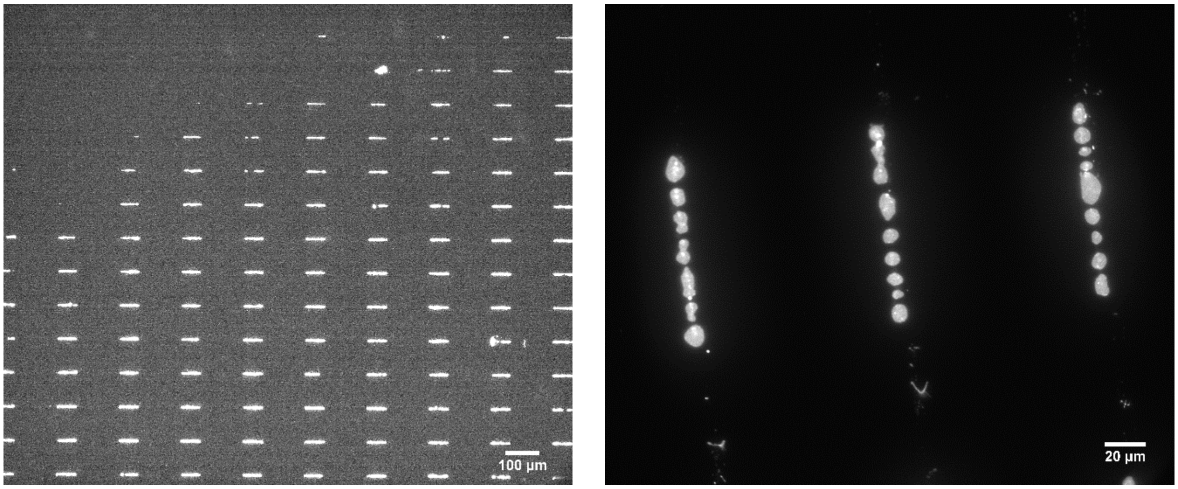

**Fig. S7:** Inconsistent micropattern quality for micropatterns micro-contact printed on carbon coated coverslips. Micro-contact printed GFP Oregon green gelatin on carbon-coated glass coverslips shows inconsistent pattern transfer (left). Non-continuous patterns (right).

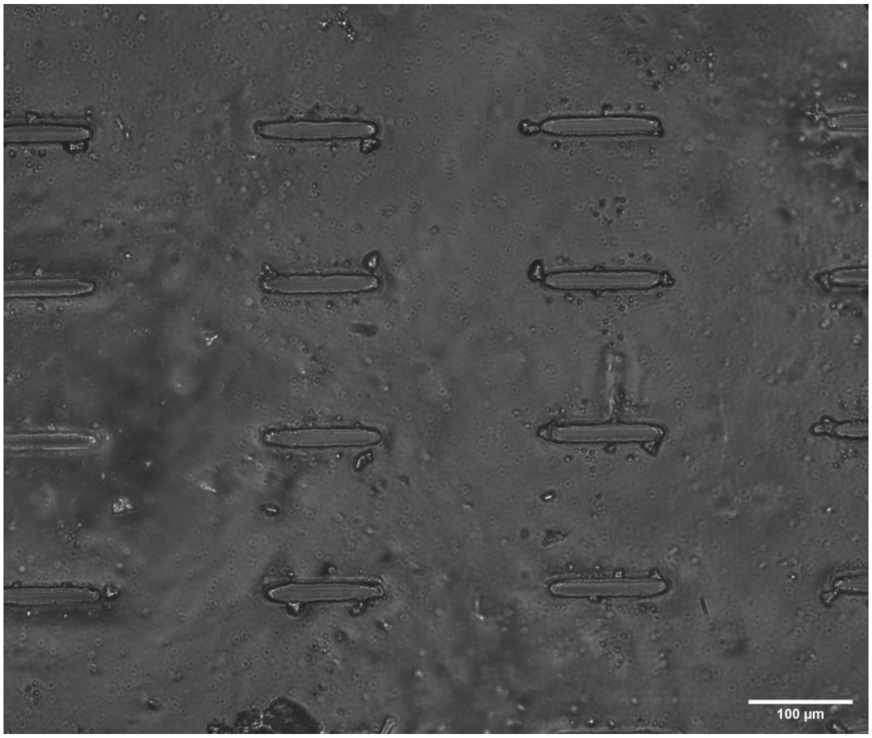

**Fig. S8:** The PDMS stamp used for micro-contact printing, post micro-contact printing. An approximation of average aspect ratio of the eight central features shown above is 1:7.6, closer to the template than the micro-contact printed features. The stamps edges appear rounded, and not sharp.

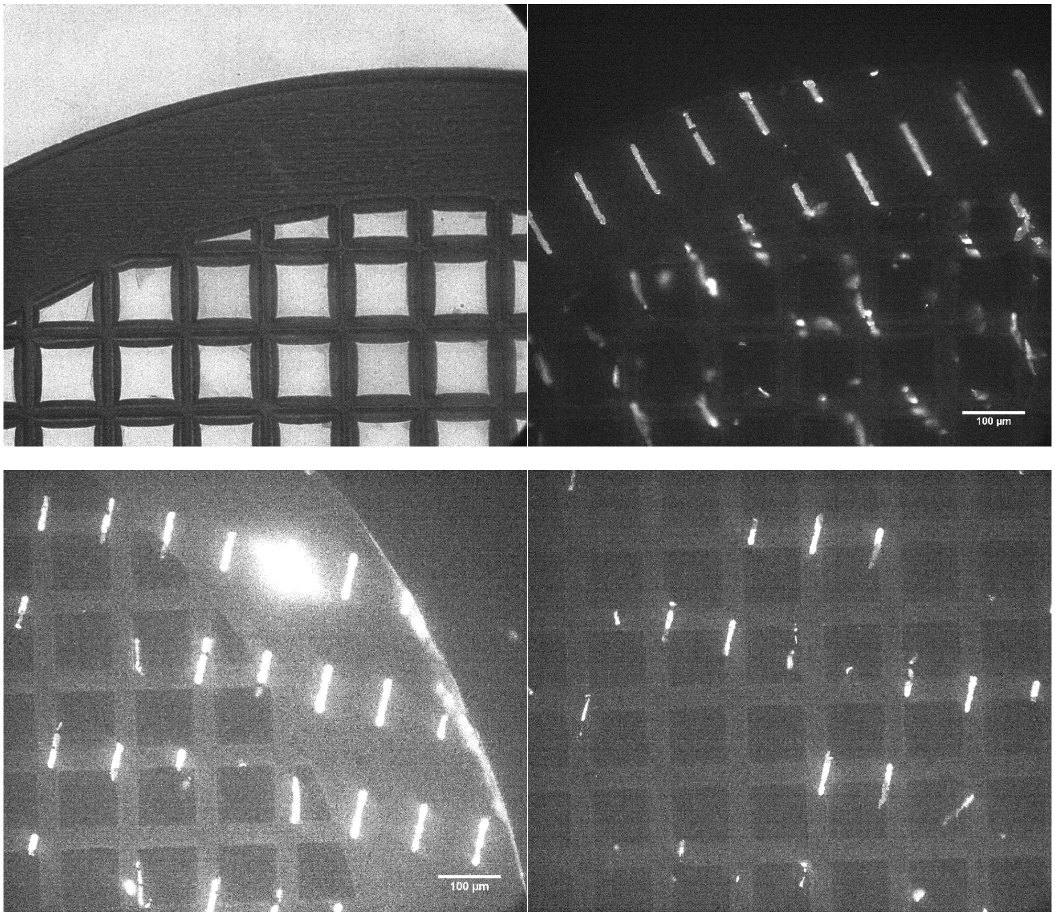

**Fig. S9**: EM grid integrity is compromised by micro-contact printing. Top left shows damaged holey carbon thin film following contact with the stamp. Top right and bottom two images show that protein was primarily deposited on areas of the grid where there is metal and not on the suspended carbon.

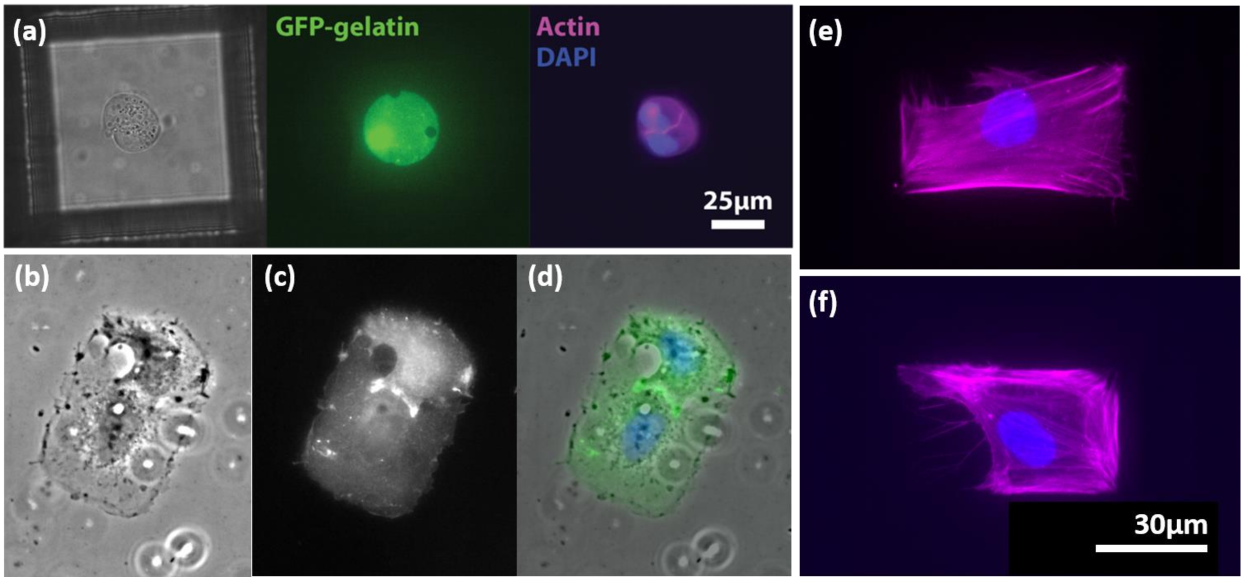

**Fig. S10**: A variety of cell types are confined to ECM micropatterns generated on carbon films via maskless photopatterning. (a) MDCKII cell doublet is confined to a circle of gelatin on the holey carbon surface of an EM grid. (b) brightfield. (c) GFP-gelatin. (d) Composite. (e) and (f) are single HFF cells on rectangular micropatterns.

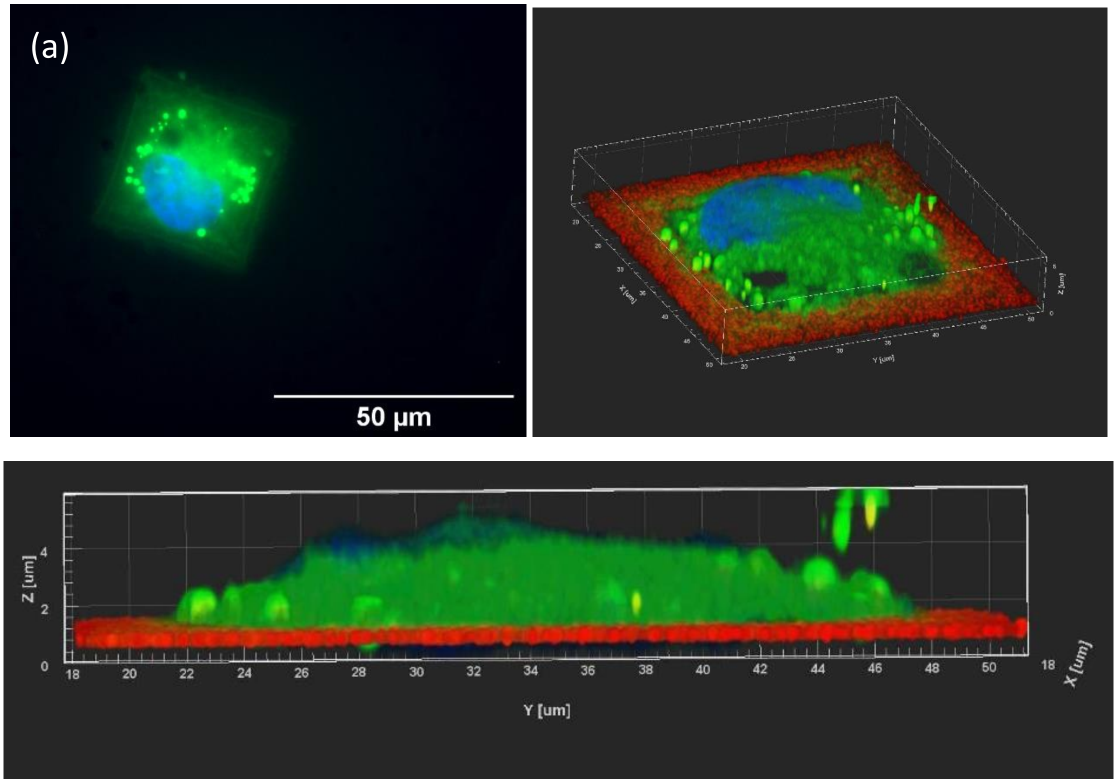

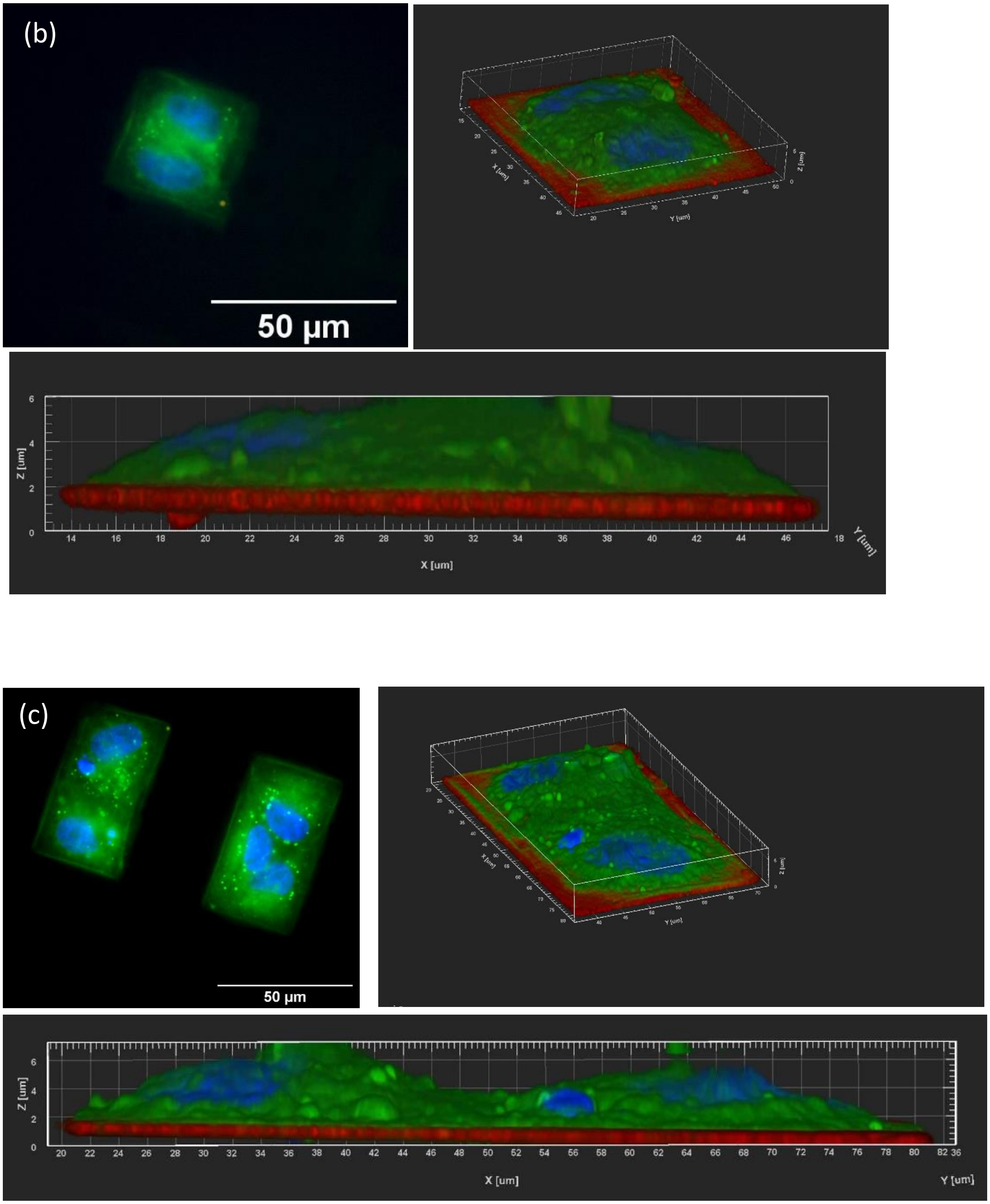

**Fig. S11:** <u>High resolution and confocal fluorescence microscopy images of MDCKII cells plated on maskless photopatterned fibronectin islands.</u> (a) A single MDCKII cell on maskless photopatterned 35x35 µm fibronectin square. (b) A cell pair on an identical sized square. (c) A cell pair on a 35x60 µm rectangle. All samples were grown on a carbon coated glass substrate and fixed 20 hours after cell seeding. Actin is labeled with green fluorescent protein and nuclei are stained with DAPI.

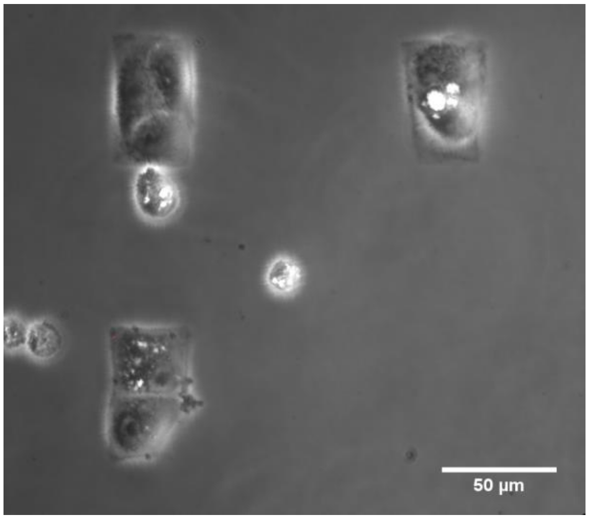

**Fig. S12:** Cell confinement to maskless photopatterned micropatterns. Phase contrast image of MDCKII cell groups confined to 35x60 µm rhodamine fibronectin rectangles on a continuous carbon film. A cell pair is confined within the bottom left of the image, while a triplet is confined in the top left.

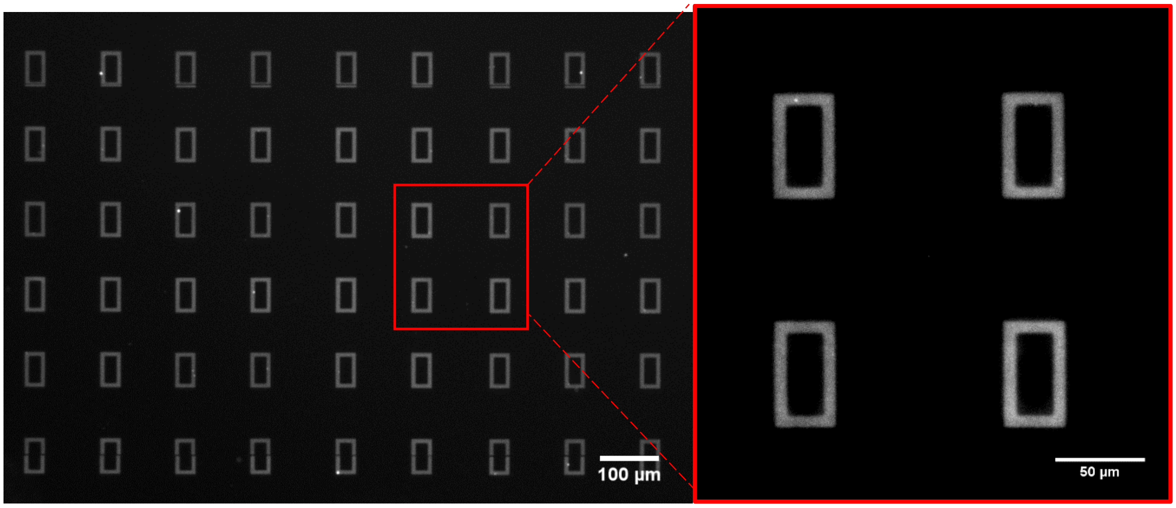

**Fig S13:** Protein micropatterns on continuous carbon film. 35x60 µm rectangles patterned on continuous carbon with 488 Oregon green gelatin (100 µg/ml) at 10x (left) and 40x (right).
